## Supplemental Materials for "Ladder shaped microfluidic system enabling rapid antibiotic susceptibility testing with standardized concentration panel"

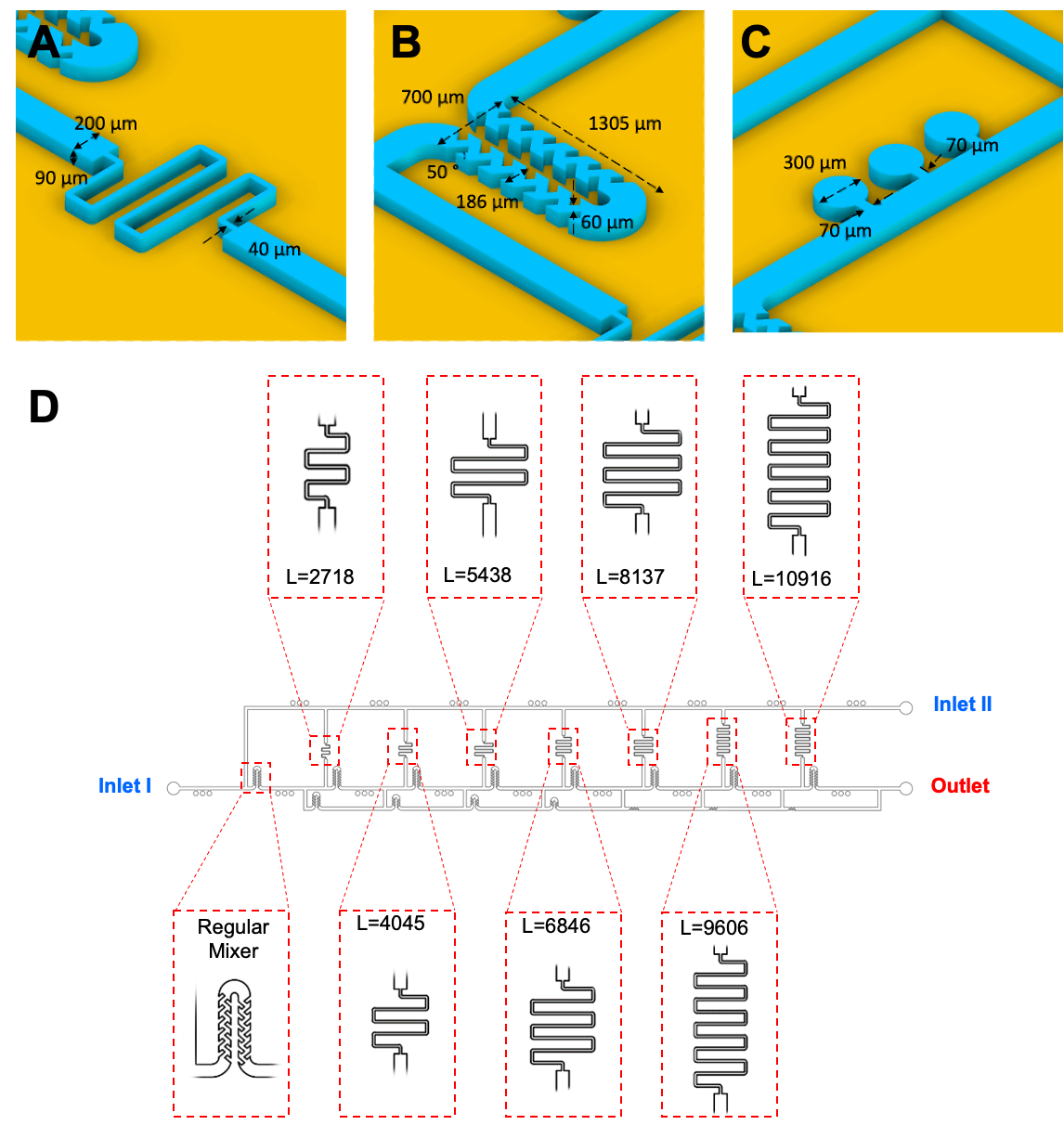

Figure S1. Detailed dimensions of the ladder shape microfluidic system

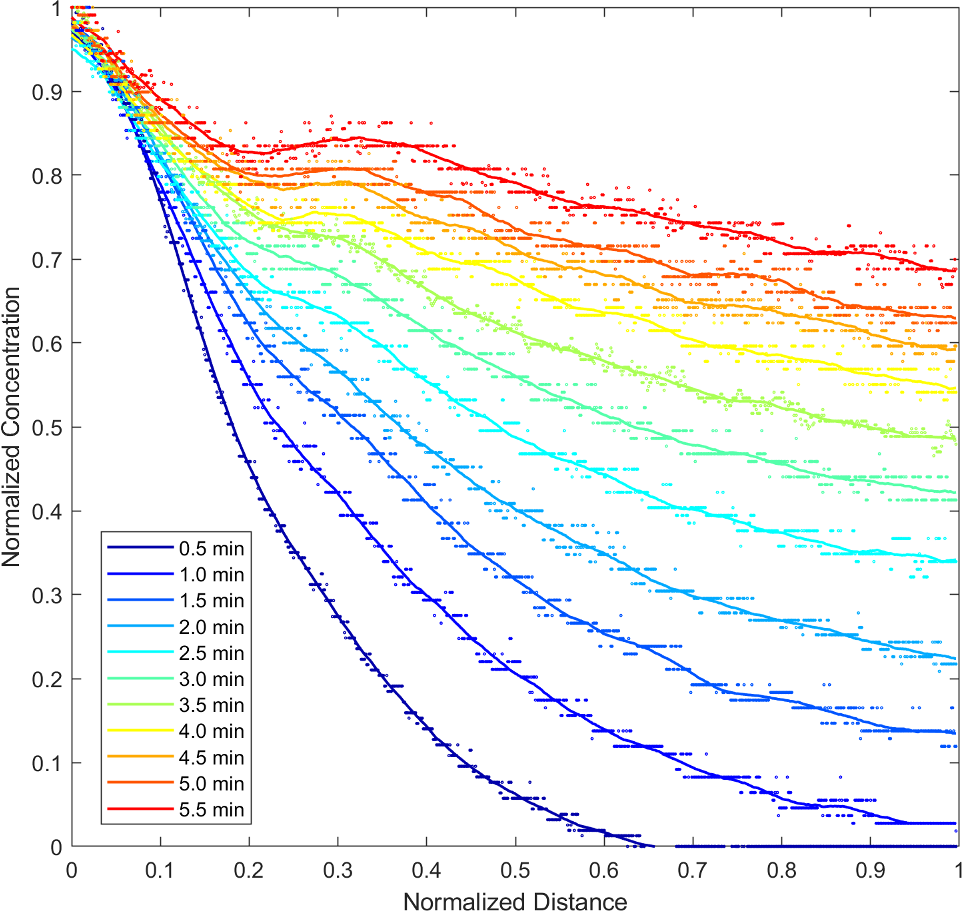

Figure S2. Relative concentration profiles showing the kinetic of resazurin diffusion into a microchamber overtime
